# The tomato yellow leaf curl virus C4 protein alters the expression of plant developmental genes correlating to leaf upward cupping phenotype in tomato

**DOI:** 10.1101/2021.09.14.460403

**Authors:** Chellappan Padmanabhan, Yi Zheng, Md Shamimuzzaman, Jennifer R. Wilson, Zhangjun Fei, Kai-Shu Ling

## Abstract

*Tomato yellow leaf curl virus* (TYLCV), a monopartite begomovirus in the family *Geminiviridae*, is efficiently transmitted by the whitefly, *Bemisia tabaci*, and causes serious economic losses to tomato crops around the world. TYLCV-infected tomato plants develop distinctive symptoms of yellowing and leaf upper cupping. In recent years, excellent progress has been made in the characterization of TYLCV C4 protein function as a pathogenetic determinant in experimental plants, including *Nicotiana benthamiana* and *Arabidopsis thaliana*. However, molecular mechanism leading to disease symptom development in natural host plant tomato has yet to be characterized. The aim of the current study was to generate transgenic tomato plants expressing the TYLCV *C4* gene and evaluate differential gene expression through comparative transcriptome analysis between the transgenic *C4* plants and the transgenic green fluorescent protein (*Gfp*) gene control plants. Transgenic tomato plants expressing the TYLCV C4 developed phenotypes, including leaf upward cupping and yellowing that are similar the disease symptom expressed on tomato plants infected with TYLCV. In a total of 241 differentially expressed genes identified in the transcriptome analysis, a series of plant development-related genes, including transcription factors, glutaredoxins, protein kinases, R-genes and microRNA target genes, were significantly altered. These results provide further evidence to support the important function of the C4 protein in begomovirus pathogenicity. These transgenic tomato plants could serve as basic genetic materials for further characterization of plant receptors that are interacting with the TYLCV C4.

## 1. Introduction

Tomato (*Solanum lycopersicum* L.) is one of the most economically important and widely grown vegetable crops in the world. Viral diseases are a major factor limiting tomato production. Tomato yellow leaf curl virus (TYLCV), a whitefly (*Bemisia tabaci*)-transmitted begomovirus, has caused serious economic losses to tomato productions worldwide [1, 2]. TYLCV, in the genus *Begomovirus* and the family *Geminiviridae*, has a monopartite genome of a single-stranded circular DNA molecule of ~2.8 kb in size. The TYLCV genome contains six open reading frames (ORFs), including two ORFs in virion (V) sense orientation, *V1* and *V2*, encoding coat protein and pre-coat, respectively, and four ORFs in complementary (C) orientation, *C1, C2, C3* and *C4*, encoding proteins responsible for virus replication, trans-activation, accumulation and induction of symptoms, respectively. Furthermore, three geminivirus-encoded proteins, C2, C4 and V2, also play a role in RNA-silencing suppression [3].

TYLCV-encoded *C4* is embedded within a larger ORF, *C1*, in a different reading frame. C4 is a relatively conserved protein which may display diverse biological functions in monopartite and bipartite geminiviruses. In monopartite begomoviruses, expression of tomato leaf curl virus (TLCV) *C4* showed virus-like symptoms in transgenic tobacco and tomato plants [4]. The C4 protein of tomato leaf curl Yunnan virus (TLCYnV) induced severe developmental abnormalities in *Nicotiana benthamiana* [5] and great progress has been achieved to identify several host factors that are interacting with the TLCYnV C4 [6, 7, 8, 9].

There are likely multi-functional roles for the TYLCV C4 that would need to be further explored [10, 11, 12]. It has been shown that the TYLCV C4 protein interacts with the BARELY ANY MERISTEM 1 (BAM1) and suppresses the cell-to-cell movement of RNAi signals [13] and chloroplast-dependent anti-viral salicylic acid (SA) biosynthesis in *Arabidopsis* [14] Another study in *Arabidopsis* demonstrated that the TYLCV C4 protein interacted broadly with plant receptor-like kinases [15] It has been suggested that due to its interaction with CLV1, C4 inhibits the cooperative interaction between CLV1 and WUSCHEL, affecting their function in maintenance of stem cells in shoot meristems, resulting in the leaf curl-like symptoms [16]. These recent development in TYLCV C4 functional studies in model plant species are very encouraging and we were aiming in characterizing the TYLCV C4 function in natural host plant tomato.

In the present study, transgenic tomato plants expressing TYLCV *C4* gene developed plant stunting, leaf upward cupping and yellowing phenotypes that are resembling those disease symptoms in tomato plants infected by TYLCV. To characterize what types of genes and metabolic pathways that are affected by expressing TYLCV *C4* gene in transgenic tomato plants, we conducted comparative transcriptome analysis and identified a series of genes encoding for transcription factors, glutaredoxins, protein kinases, R-genes and microRNAs were significantly altered.

## 2. Results

### 2.1. Development of TYLCV C4 Expressing Transgenic Tomato Plants

To develop transgenic tomato plants expressing TYLCV *C4*, a full sequence of the *C4* gene of a TYLCV isolate from Florida, USA was synthesized and cloned into the plant expression vector PEG101 (Gateway) between the cauliflower mosaic virus (CaMV) 35S promoter and nopaline synthase (NOS) terminator. Transgenic tomato plants were generated using *Agrobacterium* (LBA4404)-mediated transformation of the tomato ‘Moneymaker,’ a cultivar that is very susceptible to TYLCV infection. We initiated an *Agrobacterium* transformation with 353 explants (leaf-discs), which resulted in 28 plantlets in the selection media, from which we recovered 18 rooted plants. Among those, two transgenic tomato lines (designated C4-C1 and C4-C5) were selected for further analysis. These T_0_ and T_1_ transgenic C4 plants developed phenotypes of plant stunting, upward leaf cupping and leaf yellowing, which resembled typical tomato yellow leaf curl disease symptoms on tomato plants infected by TYLCV (Figs. 1 and 2).

**Figure 1.**
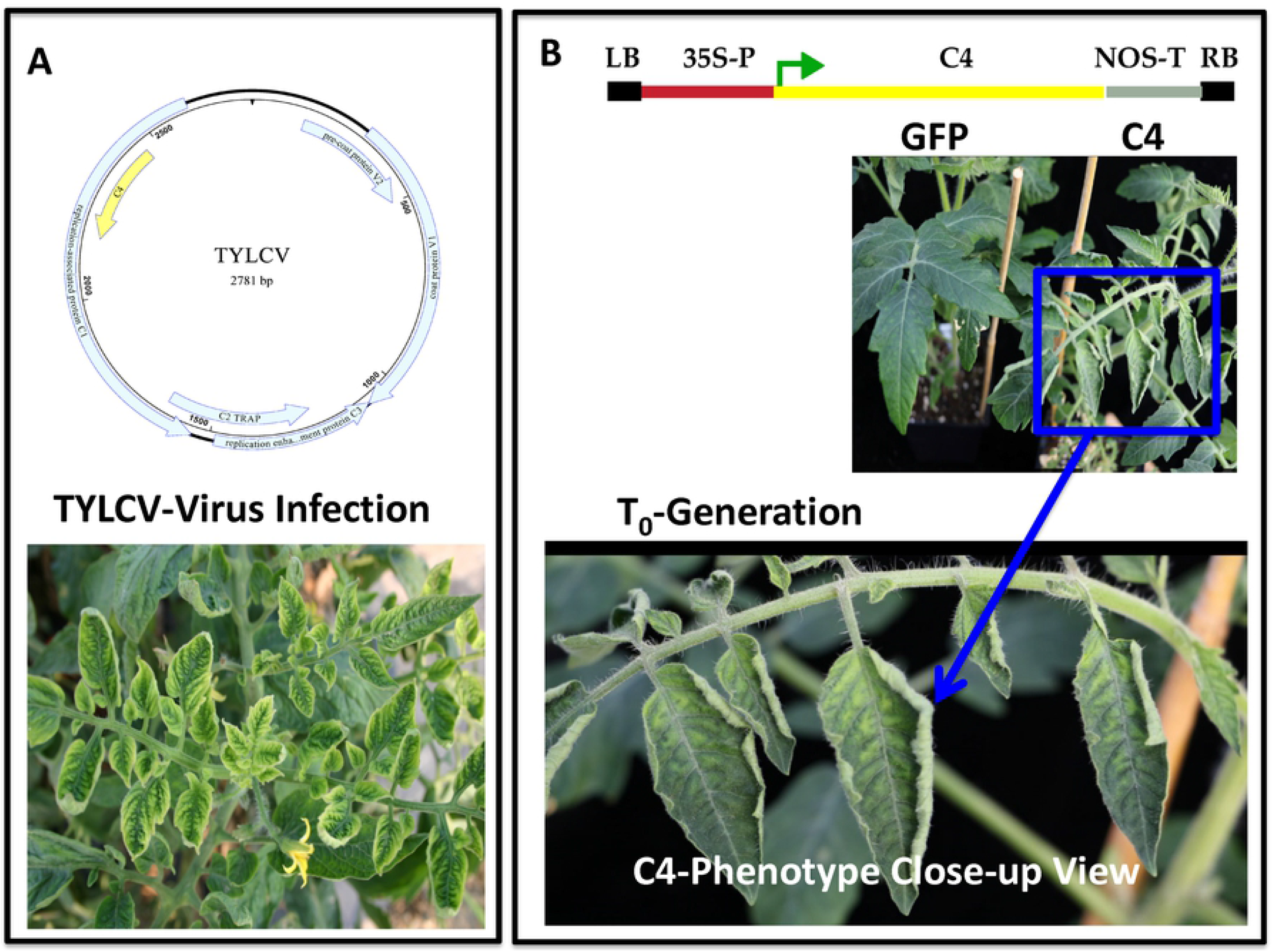
Development of transgenic tomato plants expressing the TYLCV *C4* gene. **A**) A schematic model for TYLCV (upper panel) showing its genome organization as a typical monopartite begomovirus and yellow leaf curl symptoms on tomato plants naturally infected by TYLCV (lower panel). **B**) A schematic model of the T-DNA region between the right border (RB) and Left border (LB) depicting the TYLCV *C4* gene under 35S promoter control and a NOS terminator (top panel) used to develop transgenic tomato plants. Aside-by-side comparison of the phenotypes (middle panel) displayed on a *Gfp*-transgenic plant (left side) and a TYLCV *C4*-transgenic tomato plant (right side). A close-up view of the yellow leaf curl disease-like phenotypes (yellowing and upward cupping leaves) displayed on a TYLCV *C4*-transgenic plant (lower panel).

**Figure 2.**
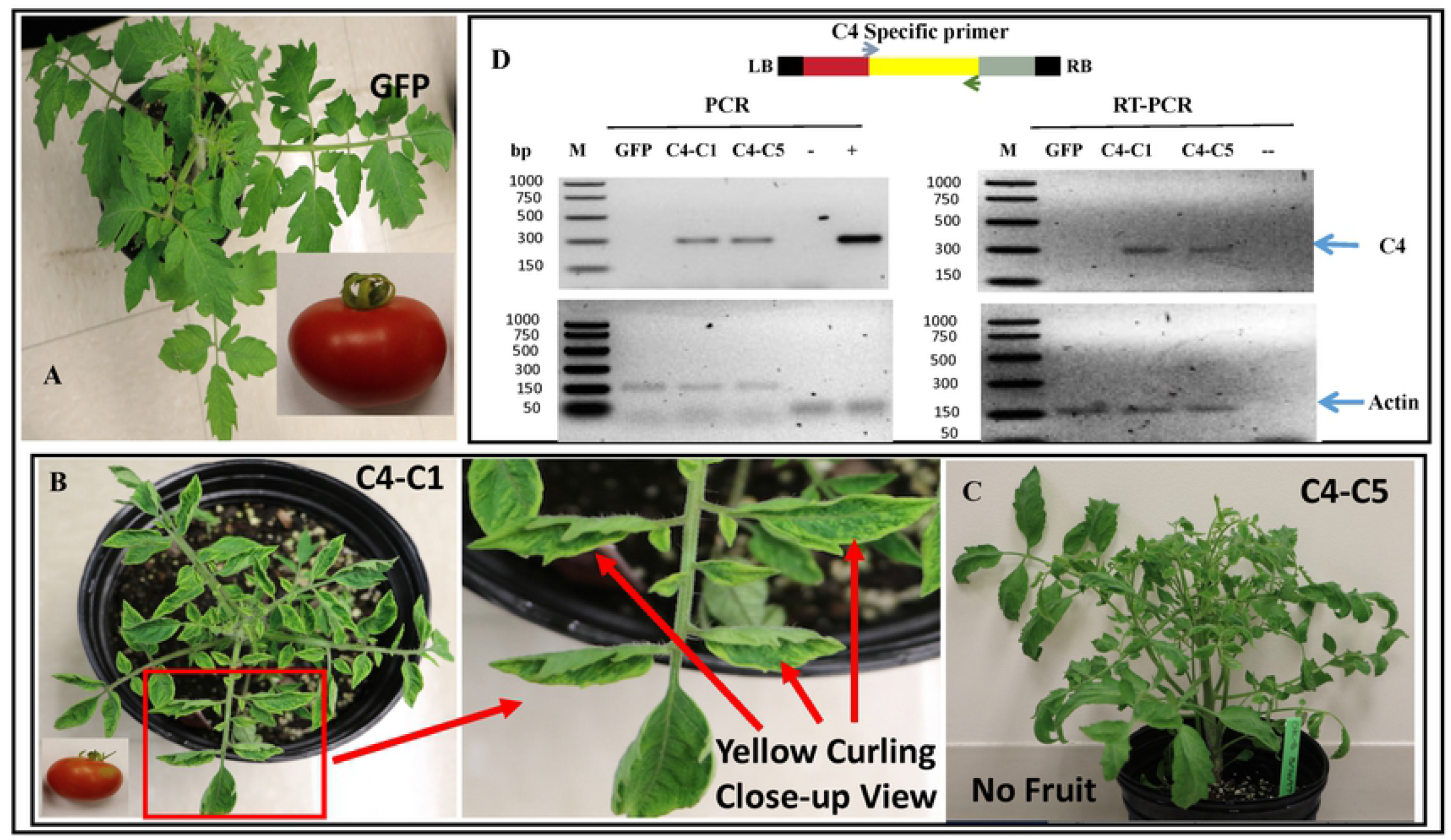
Biological and molecular characterization of the TYLCV *C4* gene expression on transgenic tomato plants. **A**) A control *Gfp*-transgenic plant with normal phenotypes in plant growth and fruit development. **B**) Transgenic tomato plants expressing TYLCV *C4* gene developed upward leaf cupping phenotypes resembling TYLCV infection on tomato, the T_0_ line ‘C4-C1’ (with a close-up view on a leaflet) was able to generate fruits, albeit of a smaller size, which allowed us to evaluate plants in the T_1_ plants. C). Another independent line ‘C4-C5’ expressed a similar leaf curl phenotype but bearing no fruit. **D**) Molecular characterization of the transgene *C4* expression in transgenic tomato plants using their respective DNA preparations with gene-specific primers (top panel) in polymerase chain reaction (PCR) (left panel) or RNA preparations by reverse transcription PCR (RT-PCR) (right panel). Two TYLCV *C4*-transgenic tomato plants (C4-C1 and C4-C5) along with a control *Gfp*-transgenic plant (GFP) were used. “+” and “-” were plasmid DNA with or without TYLCV *C4* sequence, respectively. In the bottom panels, an endogenous host gene “Actin” was used as internal quality control for DNA or RNA preparations used for their respective reactions.

The two transgenic lines induced similar phenotypes, with upward leaf cupping and plant stunting, while producing smaller size of fruits (line ‘C4-C1’) or no fruit (line ‘C4-C5’) (Fig. 2). In contrast, similarly generated control transgenic *Gfp* plants presented with normal phenotype (Fig. 2). The insertion of the transgenes *C4* or *Gfp* in those transgenic plants was validated using polymerase chain reaction (PCR) and their expression was confirmed via reverse-transcriptase (RT)-PCR with their respective gene-specific primers (Fig. 2; Supplementary Fig. S1). These analyses demonstrated that the transgenic C4 plants with the yellow leaf curl disease-like phenotype contained and expressed the expected TYLCV *C4* transgene. Observation of disease-like phenotype in the stable transgenic tomato plants offered a golden opportunity to unravel the function of the TYLCV *C4* gene. To characterize inheritance of the disease-like phenotype in the T_1_ transgenic plants, we observed a segregation of leaf curl-like phenotype in the T_1_ seedlings generated from the transgenic tomato plants expressing TYLCV *C4*. RT-PCR tests confirmed the presence of transgene expression in those T_1_ plants exhibiting plant stunting and leaf upward cupping phenotype. On the other hand, the control transgenic tomato plants expressing the green fluorescent protein (*Gfp*) gene exhibited a normal appearance phenotype as those of non-transgenic plants (Fig. 1).

### 2.2. Comparative Transcriptome Analysis of Transgenic C4 and Green Fluorescent Protein (Gfp) Control Plants

To understand the underlying molecular mechanism leading to the yellow leaf curl disease-like phenotype in the transgenic *C4* plants, we conducted a comparative transcriptome profile analysis to identify differentially expressed genes between *C4* transgenic plants and the control *Gfp* transgenic plants. Among them, three individual T_1_ transgenic plants from the ‘C4-C1’ line and three transgenic tomato plants expressing the *Gfp* at the same growth stage under the same environmental conditions in the same greenhouse were selected for transcriptome analysis. Overall, an average of ~21.5 million raw reads per library were generated. After adapter trimming and removal of low-quality reads and rRNA sequences, an average of ~17.1 million high quality clean reads were obtained, with ~15.9 million of those reads mapping to the tomato genome (version SL3.0) (Supplementary Table S1). Values of Pearson’s correlation coefficients for all biological replicates were high, suggesting highly reproducible data generated by RNA-Seq (Supplementary Table S2).

Among these RNA-seq libraries, a high number of reads were mapped to the target transgenes, 105 to 285 reads to the TYLCV *C4* and 10,834 to 20,311 reads to the *Gfp* (Table 1). We also observed a similar trend when using normalized expression of the *C4* and *Gfp* transgenes in RPKM (Reads Per Million Per Kilobase Mapped Reads) (Table 1). This provided further evidence supporting the expression of the target transgenes in their respective transgenic plants, which laid a foundation for a comprehensive analysis of global gene expression in transgenic tomato plants to examine their responses in association with expression of a disease-like phenotype in the *C4* transgenic plants.

**Table 1.** Transgene expression analysis of RNA-seq reads mapped to the *C4* or *Gfp* transgene

| Library | Genome Mapped Reads (Million) | Transgene Length (Kb) | Transgene Read Counts | Normalized Transgene Expression (RPKM)<sup>a</sup> |
| --- | --- | --- | --- | --- |
| C4-C1-1 | 15.51 | 0.298 | 105 | 22.72 |
| C4-C1-2 | 24.98 | 0.298 | 285 | 38.29 |
| C4-C1-3 | 15.87 | 0.298 | 131 | 27.70 |
| GFP1-1 | 11.78 | 0.721 | 10834 | 1275.58 |
| GFP1-2 | 15.9 | 0.721 | 20311 | 1771.74 |
| GFP1-3 | 11.82 | 0.721 | 12145 | 1425.10 |

<sup>a</sup>Reads were normalized in RPKM (Reads Per Million Per Kilobase Mapped Reads). Number of reads were divided by length of transgene in kilobases and total number of genome mapped reads in millions.

We identified a total of 241 differentially expressed genes (DEGs) (Supplementary Dataset S1), with 152 upregulated (Supplementary Dataset S2) and 89 down-regulated (Supplementary Dataset S3) in the transgenic *C4* plants compared to the transgenic *Gfp* plants (Fig. 3A). A pathway analysis of all DEGs showed that 126 pathways were altered (Supplementary Dataset S4). Gene Ontology (GO) term enrichment analysis revealed that 13 different functional categories were enriched in the DEGs (Fig. 3B), with glutaredoxin activity, arsenate reductase activity and cell redox homeostasis being the top three categories. Among 152 up-regulated genes, the most prominent annotation group was glutaredoxins. Among 89 down-regulated genes, the most prominent annotation group was receptor-like protein kinases (Table 2).

**Figure 3.**
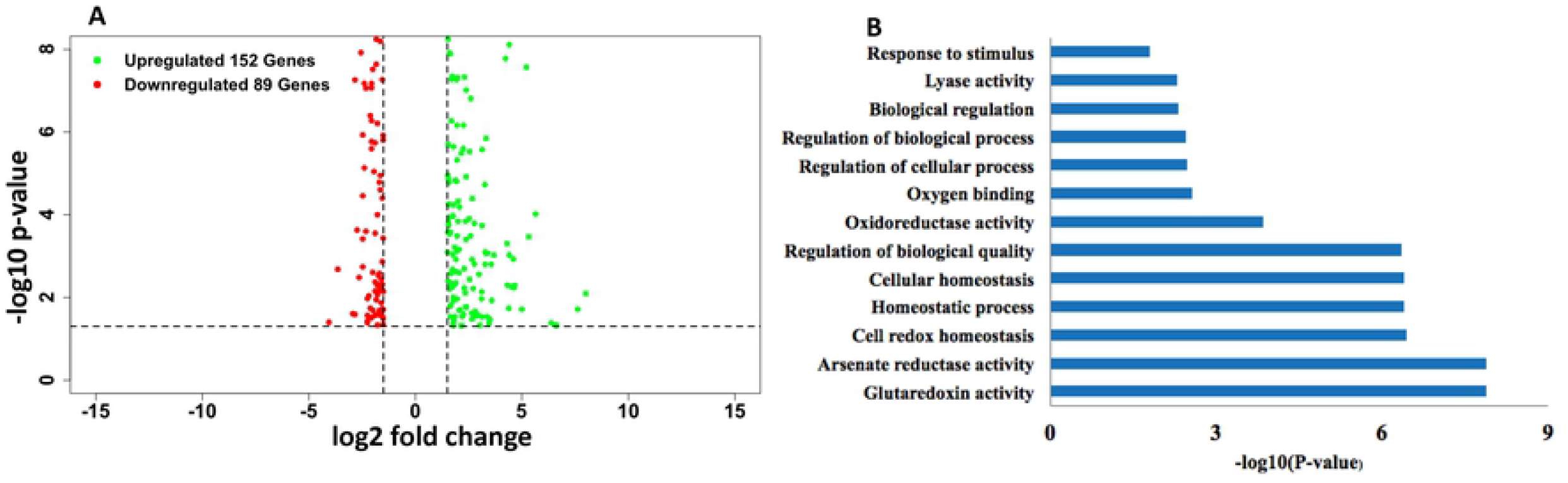
Comparative transcriptome analysis on differential gene expression between the TYLCV *C4*-transgenic tomato plants and those control transgenic plants expressing the *Gfp* gene under the same genetic background. **A**) A volcano plot showed a distribution pattern of differentially expressed genes (DEGs) with number of up-regulated (in green) or down-regulated (in red) genes in the *C4*-transgenic tomato plants over that of the *Gfp*-transgenic plants. X-axis represents -log10 (p-value) and y-axis represents log2 (fold change). Black horizontal dotted lines show the p-value cut off at 0.05. Black vertical dotted lines were drawn using log2 (fold change) cut off at −1.5 and 1.5. **B**) Gene Ontology (GO) enrichment analysis revealed 13 enriched categories of the identified DEGs, with category in the y-axis and –log10 (p-value) in the x-axis.

**Table 2.** Classification of differentially expressed genes to prominent annotation groups

| Up-regulated Genes Annotation Group | Number of Genes (152) |
| --- | --- |
| Glutaredoxin | 12 |
| Avr9/Cf-9 rapidly elicited protein | 3 |
| Cytochrome P450 | 3 |
| Late embryogenesis abundant family protein | 3 |
| MADS box transcription factor | 3 |
| MYB transcription factor | 3 |
| Plant-specific domain TIGR01589 family protein | 3 |
| WRKY transcription factor | 3 |
| Unknown Protein | 30 |
| Others | 89 |
| Down-regulated Genes Annotation Group | Number of Genes (89) |
| Receptor like protein kinase | 4 |
| Xyloglucan endotransglucosylase/hydrolase | 3 |
| Cytochrome P450 | 3 |
| bZIP/bHLH transcription factor | 3 |
| Unknown Protein | 5 |
| Others | 71 |

### 2.3. Characterization of Selected Differentially Expressed Genes

Further classification placed DEGs into different regulatory groups such as transcription factors, protein kinases, R-genes, and microRNA target genes. From the GO enrichment analysis, we determined that the oxidoreductase activity of glutaredoxin (GRX) was one of the most highly enriched categories of DEGs (Fig. 3B). GRXs allow for redox regulation of protein activity by reversibly glutathionylating or reducing disulfide bridges in their targets and plant developmental function (Table 3). Twelve glutaredoxin genes were differentially expressed and all of them were induced in the *C4* transgenic plants (Table 3).

**Table 3.** Glutaredoxin genes differentially expressed between the transgenic *C4* plants and the transgenic *Gfp* control plants

| Gene ID | Annotation | log2fold | Adjusted P-value | General Functions |
| --- | --- | --- | --- | --- |
| Solyc05g051720 | Glutaredoxin | 7.99 | 0.008 | Flower development, Salicylic acid signaling, Oxidative stress, Petal development, Anther development, Floral organ primordium formation, Root development, Dwarf phenotype, and Embryo development. |
| Solyc04g011860 | Glutaredoxin | 4.38 | 0.0183 |  |
| Solyc04g011880 | Glutaredoxin | 1.63 | 0.0163 |  |
| Solyc01g067440 | Glutaredoxin | 2.75 | 1.05E-09 |  |
| Solyc04g011840 | Glutaredoxin | 7.62 | 0.0192 |  |
| Solyc05g051730 | Glutaredoxin | 3.68 | 0.0009 |  |
| Solyc04g053110 | Glutaredoxin | 3.27 | 0.0007 |  |
| Solyc04g011800 | Glutaredoxin | 3.12 | 0.0293 |  |
| Solyc04g011830 | Glutaredoxin | 4.6 | 0.0011 |  |
| Solyc06g054570 | Glutaredoxin | 4.23 | 1.66E-08 |  |
| Solyc01g067460 | Glutaredoxin | 1.95 | 6.83E-07 |  |
| Solyc09g074590 | Glutaredoxin | 2.18 | 0.0478 |  |

On the other hand, a total of 18 transcription factor (TF) genes belonging to eight different families exhibited differential expression patterns between the transgenic *C4* plants and the control *Gfp* transgenic plants, among which 14 were up-regulated while four were down-regulated in the *C4* transgenic plants (Table 4). The 14 up-regulated TFs included one basic helix-loop-helix (bHLH), two HD-ZIP, three MADS box, three MYB, one NAM/NAC, three WRKY and one LOB TF gene. On the other hand, one bHLH, two bZIP and one MYB TF gene were down-regulated.

**Table 4.** Differentially expressed genes representing transcription factors between transgenic *C4* plants and the *Gfp* control plants

| Gene ID | Annotation | Function | log2fold | Adj P-val |
| --- | --- | --- | --- | --- |
| **bHLH Family** |  |  |  |  |
| Solyc02g091690 | bHLH transcription factor | Erect leaf phenotype dwarfism | -2.4 | 7.00E-06 |
| Solyc03g006910 | bHLH transcription factor | Erect leaf phenotype dwarfism | 3.27 | 0.0015 |
| **bZIP Family** |  |  |  |  |
| Solyc07g053450 | bZIP transcription factor | Leaf cell number and cell size | -2.06 | 0.0307 |
| Solyc12g010800 | bZIP transcription factor | Leaf cell number and cell size | -1.64 | 6.00E-09 |

| **HD-ZIP Family** |  |  |  |  |
| --- | --- | --- | --- | --- |
| Solyc01g096320 | HD-ZIP transcription factor | Adaxialized leaf (upward leaf cupping) | 3.1 | 5.16E-28 |
| Solyc06g053220 | HD-ZIP transcription factor | Adaxialized leaf (upward leaf cupping) | 2.2 | 0.0288 |
| **LOB Family** |  |  |  |  |
| Solyc04g077990 | LOB domain transcription factor | Leaf primordia development | 1.62 | 0.0061 |
| **MADS Family** |  |  |  |  |
| Solyc02g065730 | MADS box transcription factor | Leaf morphogenesis | 1.73 | 4.49E-08 |
| Solyc05g056620 | MADS box transcription factor | Leaf morphogenesis | 2.37 | 0.0003 |
| Solyc02g071730 | MADS-box transcription factor | Leaf morphogenesis | 2.59 | 0.0003 |
| **MYB Family** |  |  |  |  |
| Solyc01g010910 | MYB transcription factor | Maintenance of leaf morphogenesis | -1.51 | 0.007 |
| Solyc05g008250 | MYB transcription factor | Maintenance of leaf morphogenesis | 1.79 | 0.048 |
| Solyc11g073120 | MYB transcription factor | Maintenance of leaf morphogenesis | 1.55 | 0.0001 |
| Solyc01g109670 | MYB transcription factor | Maintenance of leaf morphogenesis | 4.18 | 2.00E-10 |
| **NAM Family** |  |  |  |  |
| Solyc12g013620 | NAM/NAC transcription factor | Specification of leaflet boundaries | 1.77 | 0.0001 |
| **WRKY Family** |  |  |  |  |
| Solyc08g062490 | WRKY transcription factor | Flag leaf growth and host defense | 1.78 | 0.0119 |
| Solyc03g116890 | WRKY transcription factor | Flag leaf growth and host defense | 2.05 | 0.0203 |
| Solyc09g014990 | WRKY-like transcription factor | Flag leaf growth and host defense | 2.05 | 0.0025 |

A total of seven DEGs coding for protein kinases were identified in the RNA-seq dataset, among which three were induced and four suppressed in the transgenic *C4* plants (Table 5). Specifically, a CBL-interacting protein kinase, a calcium-dependent protein kinase and an LRR receptor-like serine/threonine-protein kinase were induced by1.5 to 2.4 log2fold. On the other hand, expression of four other protein kinase genes in the families of RLK-Pelle_LRR-XI-1, RLK-Pelle_PERK-2, RLK-Pelle_RLCK-VIIa-1 and RLK, were suppressed in the *C4* transgenic plants (Table 5).

**Table 5.** Differentially expressed genes in protein kinase families between the transgenic *C4* plants and the control *Gfp* plants

| Gene_ID | Annotation | Family | log2fold | Adj P-value |
| --- | --- | --- | --- | --- |
| Solyc08g067310 | CBL-interacting protein kinase 6 | CAMK_CAMKL-CHK1 | 2.25 | 0.0255 |
| Solyc12g099790 | Calcium-dependent protein kinase 17 | CAMK_CDPK | 1.51 | 0.0008 |
| Solyc05g007230 | Receptor like kinase, RLK | RLK-Pelle_LRR-XI-1 | -2.83 | 0.0255 |
| Solyc04g079690 | Receptor-like protein kinase 2 | RLK-Pelle_PERK-2 | -1.64 | 0.0268 |
| Solyc10g084390 | Receptor protein kinase-like protein | RLK-Pelle_RLCK-VIIa-1 | -1.88 | 0.02693 |
| Solyc11g016930 | LRR receptor serine/threonine kinase | RLP | 2.38 | 0.00001 |
| Solyc09g015840 | Receptor-like kinase | RLK | -1.51 | 0.01957 |

In addition, one gene encoding gibberellin 2-beta-dioxygenase 7 in the gibberellin (GA) biosynthesis pathway was induced by more than 2 log2fold in transgenic *C4* plants (Supplementary dataset S2). Furthermore, we identified two un-annotated microRNAs (M00148 and M00188), targeting the same gene Solyc10g007080, which encodes an Aberrant lateral root formation 5 protein, resulting in down-regulated expression (−2.94) in the transgenic *C4* plants (Supplementary dataset S3). Two different microRNAs regulating the expression of the same host gene (Solyc10g007080) is an important discovery, although their functions in regulating aberrant lateral root formation and its causal effect on plant stunting would need further study.

### 2.4. Validation of gene expression using quantitative reverse transcription PCR (qRT-PCR)

Differential expression of 14 randomly selected DEGs from the transcriptome study were validated by qRT-PCR. All genes tested by qRT-PCR were in full agreement with the expression pattern (upregulation or downregulation) observed in the RNA-seq dataset (Table 6). For all but one of these genes (Solyc11g073120), the differential expression observed via qRT-PCR was also statistically significant (p < 0.05).

**Table 6.** Summary of qRT-PCR validation of selected differentially expressed genes

| Gene ID | Annotation | RNA-seq |  | qRT_PCR |  |
| --- | --- | --- | --- | --- | --- |
|  |  | log2fold | P-value | log2fold | Adj P-value |
| Solyc04g011880 | Glutaredoxin | 1.63 | 0.0163 | 0.54 | 8.82E-06 |
| Solyc06g054570 | Glutaredoxin | 4.23 | 1.66E-08 | 2.44 | 1.01E-07 |
| Solyc01g067460 | Glutaredoxin | 1.95 | 6.83E-07 | 1.22 | 1.44E-05 |
| Solyc02g091690 | bHLH transcription factor | -2.40 | 7.00E-06 | -2.30 | 0.0052 |
| Solyc07g053450 | bZIP transcription factor | -2.06 | 0.0307 | -1.26 | 0.0354 |
| Solyc01g096320 | HD-ZIP transcription factor | 3.10 | 5.16E-28 | 0.90 | 0.0145 |
| Solyc02g071730 | MADS box transcription factor | 2.59 | 0.0030 | 2.79 | 7.67E-06 |
| Solyc03g116890 | WRKY transcription factor | 2.05 | 0.0203 | 1.18 | 0.0083 |
| Solyc01g010910 | MYB transcription factor | -1.51 | 0.0073 | -2.83 | 3.10E-09 |
| Solyc11g073120 | MYB transcription factor | 1.55 | 1.52E-04 | 0.23 | 0.1895 |
| Solyc05g007230 | Receptor-like kinase, RLK | -2.83 | 0.0255 | -1.22 | 0.0064 |
| Solyc09g014990.2.1 | WRKY-like transcription factor | -0.44 | 0.8536 | 2.05 | 0.0025 |
| Solyc03g006910.2.1 | bHLH transcription factor | -0.18 | 0.6571 | 3.27 | 0.0015 |
| Solyc08g067310.1.1 | CBL-interaction protein kinase 6 | -0.11 | 0.4041 | 2.25 | 0.0255 |

## 3. Discussion

Using stable transformed tomato plants and comparative transcriptome analysis, we were able to profile the global effects on gene expression in transgenic tomato plants expressing the TYLCV *C4* gene in comparison to the same genetic background tomato plants transformed with the *Gfp* gene. Transgenic tomato plants expressing TYLCV *C4* developed plant stunting, upward leaf cupping, and small fruit size phenotypes that resemble the yellow leaf curl disease symptoms on tomato plants naturally infected with TYLCV. Through comprehensive transcriptome profile analysis between the *C4* transgenic plants and the control *Gfp* transgenic plants, we identified a total of 241 differentially expressed genes (152 up-regulated and 89 down-regulated) using robust statistical analysis on three biologically replicated RNA-Seq with a stringent cutoff [adjusted p values < 0.05 and log2(fold change) ≥ 1.5]. We believe that these DEG analyses are highly reliable as the validation test on selected 14 genes using qRT-PCR agreed with the expression pattern generated in RNA-seq datasets used for transcriptome analysis. Our results are in agreement with several other studies which have also demonstrated the high correlation between RNA-Seq and qRT-PCR [17, 18].

Among the differentially expressed genes (DEGs) identified in our study are a series of glutaredoxins, protein kinases, transcriptions factors, and microRNAs target genes that are potentially involved in leaf tissue formation and plant development that could potentially contribute to the yellow leaf curl disease-like symptom development in transgenic tomato plants expressing the TYLCV *C4*. The result from the present study offers another evidence to support the C4 as a pathogenicity determinant for TYLCV, one of the most important tomato viruses. Although several studies have demonstrated that the C4 protein of geminiviruses is responsible for developing disease-like symptoms in tobacco, tomato, and *N. benthamiana* [4, 5]. The C4 protein has also been shown to be the pathogenicity determinant for numerous viruses in the *Geminiviridae* [4, 19, 20, 21]. Its function in TYLCV has received great attention in recent years using model plants, *Arabidopsis* and *N. benthamiana* in their studies [12, 14, 18, 22,]. Previously, Rojas and colleagues showed that TYLCV C4 is localized to the cell periphery, thus suggesting it may be involved in mediating virus cell-to-cell movement [23]. However, as the *C4* gene is totally embedded inside the *C1* open reading frame in TYLCV, this cell-to-cell movement may be attributed to the C1 protein’s function as evidenced in other bipartite begomoviruses. Another study [24] suggested that the TYLCV *C4* protein is likely a pathogenicity factor due to its interaction with and suppression by a host resistance factor to restrict virus systemic movement.

We identified a total of seven DEGs in the protein kinase families, four of which are receptor-like kinases (in the families of RLK-Pelle_LRR-XI-1, RLK-Pelle_PERK-2, RLK-Pelle_RLCK-VIIa-1, and RLK), and all are down-regulated (Table 5). Geminivirus-encoded C4/AC4 proteins have previously been shown to interact with RLKs, including CLV1 in the CLAVATA 1 (CLV1) clade [16, 22, 25], as well as BAM1 and BAM 2 [26, 27]. The targeting of BAM1 and BAM2 by TYLCV C4 has been shown to block RNAi signal spread from cell to cell [13]. In addition to two RLKs (BAM1 and BAM2) that have previously been shown to be involved in TYLCV C4 functions [12, 13, 14, 22], our transcriptome analysis also revealed the suppression of four RLK genes in the transgenic *C4* tomato plants, indicating that the RLK-mediated plant defense system may have been compromised, leading to the development of a disease-like phenotype in the transgenic tomato plants. Thus, these four RLKs identified in the present study deserve further characterization on their functions in relationship to the TYLCV resistance and susceptibility in tomato plants.

We identified a total of 12 glutaredoxins (GRXs, also known as thioltransferases) that were induced in the *C4* transgenic tomato plants, with all of them being up-regulated. GRXs are small redox enzymes of approximately one hundred amino acid residues that use glutathione as a cofactor [28]. In plants, GRXs are involved in flower development and salicylic acid signaling [29], and GRXs are well-documented to be involved in oxidative stress responses [29]. Studies revealed that two members of a land plant-specific class of GRXs, ROXY1 and ROXY2, are required for petal development in *Arabidopsis* [30]. Further studies revealed that ROXY1 interacts with several TGA transcription factors, including TGA2, TGA3, TGA7, and PERIANTHIA (PAN); the function of PAN is floral organ primordium formation [31] and root development [32], thus supporting the role of GRXs in these processes. Overexpression of a rice glutaredoxin (OsGRX6), affects hormone and nitrogen status in rice plants, resulting in a dwarf phenotype [33] whereas overexpression of OsGrxC2.2 resulted in abnormal embryos and an increased grain weight in rice [34]. In our study, we observed a stunting (dwarf) phenotype in the *C4*-transgenic plants (Fig. 2), suggesting that C4 may play a role in plant development by interfering with hormone and nitrogen status, similar to the effects of overexpressing *OsGRX6* in rice [33].

Expression of a series of leaf development transcription factors (TFs), including those in the bHLH, bZIP, HD-ZIP, NAC/NAM, MADS box, LOB, MYB and WRKY families, were altered in the *C4*-transgenic plants (Table 4). These leaf development transcription factors could be involved in functions such as regulating leaflet boundary, leaf primordial development, leaf morphogenesis, and leaf cell number and size, which may potentially lead to the leaf upward cupping phenotype.

The bHLH transcription factors, one of the largest TF super-families in plants, can participate in a broad range of growth and developmental signaling pathways. In the transgenic *C4* plants, two bHLH TFs were differentially expressed: one induced and another suppressed. Plant bHLH proteins have the potential to be involved in regulating a multiplicity of transcriptional programs. Experimental evidence reveals that bHLH genes make a significant contribution to the specification of stomata in plants [35]. On the other hand, HLH/bHLH transcription factors could have an opposite effect in mediating brassinosteroid regulation of cell elongation and plant development, and their overexpression resulted in an erect leaf phenotype in rice and dwarfism in *Arabidopsis* [36]. In another study, Ichihashi and colleagues [37] demonstrated that the bHLH transcription factor SPATULA controls final leaf size in *Arabidopsis*.

Next, some of the altered TF genes in the *C4*-transgenic plants belong to the bZIP family. TYLCV *C4* mediated a strong suppression of two bZIP genes, which may ultimately alter normal plant development, resulting in an enhanced disease-like leaf curl phenotype in the *C4*-transgenic tomato plants. bZIP TFs play crucial roles in plant development, signaling and responses to abiotic/biotic stimuli, including abscisic acid (ABA) signaling, hypoxia, drought, high salinity, cold stress, hormone signaling, light responses, osmotic stresses and pathogen defense [38, 39].

In contrast to the suppression of bZIP TFs, two TFs in the homodomain-leucine zipper (HD-ZIP) family were induced in the transgenic *C4* plants. [40] demonstrated that loss-of-function mutations in two HD-ZIPII transcription factors (athb4 and hat3) resulted in severely abaxialized and entirely radialized leaves. Conversely, overexpression of HAT3 results in adaxialized leaf development. Our data agree with the second aforementioned study as the overexpression of two HD-ZIP TFs is correlated with adaxialized leaf development (upward leaf cupping) in the transgenic *C4* tomato plants.

The NAC transcription factors, including <u>N</u>AM (no apical meristem), <u>A</u>TAF (Arabidopsis transcription activation factor), and <u>C</u>UC (cup-shaped cotyledon), have a conserved NAC domain (derived from the first letter of each gene). The transgenic *C4* tomato plants with abnormal upward leaf cupping phenotype also had an elevated expression on one of the NAC domain transcription factors. The NAC proteins are thought to be involved in developmental processes, including formation of the shoot apical meristem (SAM), floral organs, and lateral shoots [41]. Two independent studies have also provided evidence for microRNA-mediated regulation of CUC1 [42] and CUC2 [43].

The MADS-box transcription factors are important regulators of plant developmental pathway genes. Our study determined that expression of three MADS box TF genes were induced in the *C4*-transgenic plants, implicating their involvement in flower development. Previous studies have shown that members of the MADS-box family are known to be involved predominantly in developmental processes, including flowering time, floral meristem identity, floral organogenesis, fruit formation, seed pigmentation and endothelium development [44, 45].

We observed an up-regulation of one transcription factor in the LOB family. LOB TFs play important functions in maintaining lateral organ boundaries [46]. For example, the rice OsAS2 gene, a member of the LOB domain family, functions in regulating shoot differentiation and leaf development. Transgenic plants overexpressing the OsAS2 gene showed aberrant twisted leaves [47]. It is reasonable to speculate that the increased expression of LOB contributes to the development of leaf upward curling phenotype in the *C4*-transgenic tomato plants.

We also observed that four transcription factors in the MYB family were altered in the present study. One was suppressed, and three others induced in the transgenic *C4* plants. It is possible that alternation in the expression of these MYB genes led to the adverse effect on flower and fruit production and development as observed in the transgenic *C4* tomato plants. The MYB family is a part of a large family of transcription factors found in plants and animals. The MYB TFs are regulators of many plant processes, including responses to biotic and abiotic stresses, development, differentiation, metabolism, and defense [48, 49].

Finally, modulated expression of three WRKY TF genes in the transgenic *C4* tomato plants may lead to suppression of the host defense to TYLCV infection. The WRKY family transcription factors are key regulators of many processes in plants, including biotic and abiotic stresses, seed dormancy and germination, and other developmental process [50, 51]. It has been reported that AtWRKY52 contains a TIR–NBS–LRR (Toll/interleukin-1 receptor–nucleotide-binding site-leucine-rich repeat) domain acts together with RPS4 to provide resistance against fungal pathogen *Colletotrichum higginsianum* and bacterial pathogen *Pseudomonas syringae* [52].

## 4. Conclusions

A comprehensive understanding of key host genes involved in plant response to virus infection is a fundamental knowledge in developing an effective strategy for disease management. Transgenic tomato plants expressing the *C4* gene of TYLCV developed an upward leaf cupping phenotype that resembles the yellow leaf-curl disease symptoms on tomato plants infected by TYLCV, indicating importance of the C4 protein of TYLCV (Fig. 4). Through comparative transcriptome analysis between the *C4*-transgenic plants and the control *Gfp*-transgenic plants, a series of differentially expressed genes and their regulatory networks were uncovered. In the case of *C4*-transgenic tomato plants showing a leaf upward cupping phenotype, the expression of a series of important transcription factor family genes were altered. Our analysis revealed that the C4 protein of TYLCV interferes with the expression of several transcription pathway genes, potentially leading to the leaf upward cupping phenotype (Fig. 4). A basic understanding of this virus-encoded virulence factor and associated host responses on the molecular level is important for viral disease management.

**Figure 4.**
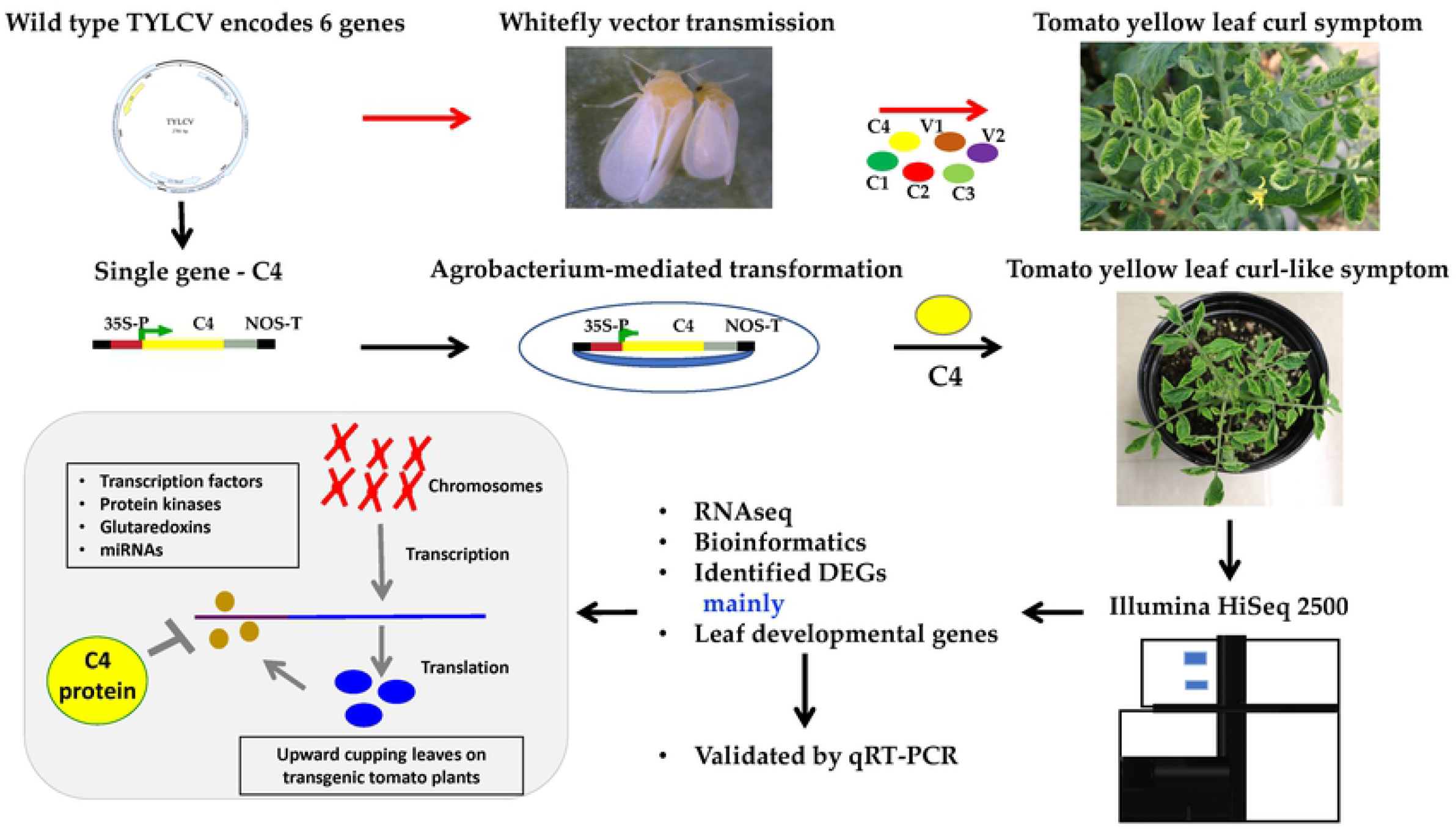
A schematic flow chart depicts the potential functional interference of the TYLCV C4 protein to a series of plant developmental genes, especially those involving in transcriptional regulation, protein kinase, Glutaredoxin and gene silencing pathways. The top panel shows a natural field infection of tomato plants by TYLCV through transmission by viruliferous whiteflies. The middle and lower panels showed key steps in the development of transgenic tomato plants expressing the TYLCV *C4* gene, transcriptome analysis and predicted functional interference on host genes that are regulating plant development, resulting in yellow leaf curl disease-like phenotypes.

## 5. Material and Methods

### 5.1. Generation of Binary TYLCV C4 Constructs

The *C4* gene of the TYLCV isolate from Florida, USA (GenBank Accession No. AY530931.1) was synthesized by IDT (Coralville, IA). The synthetic *C4* gene (C-terminus fusion) was inserted into a pENTR D TOPO vector and transformed into Top 10 chemically competent cells (Invitrogen).

Colonies were selected on kanamycin containing LB plates and the cloned *C4* sequence was confirmed using Sanger sequencing. A positive clone was recombined with a plant expression vector, pEG101, using LR clonase (Invitrogen, USA) to insert the TYLCV *C4* gene in between the CaMV 35S promoter and nopaline synthase (NOS) terminator. The sequence confirmed *C4* gene in the pEG101 background was mobilized into *Agrobacterium tumefaciens* strain LBA4404 by electroporation. Agrobacterium colonies selected on a YM agar plate containing kanamycin and streptomycin were used for plant transformation.

### 5.2. Tomato Transformation and Confirmation

Tomato transformation was conducted using tomato ‘Moneymaker’ following the outlined procedures [53]. The primary transformant plants were confirmed to contain the TYLCV *C4* sequence by PCR using the following primer pair: KL14-390 C4N-1F: 5’-CACCATGGGGAACCACATCTCCAT-3’ and KL14-391 C4N-1R: 5’-TTAATATATTGAGGGCCTCGGATTT-3’. As an experimental control, transgenic tomato plants with the same genetic background ‘Moneymaker’ containing the green fluorescent protein gene (*Gfp*) was previously developed [54].

For the control *Gfp*-transgenic plants, a confirmation test was conducted using the primer pair KL14-414 GFP-1F: 5’-CACCATGGGCAAGGGCGAGGAACT-3’ and KL14-415 GFP-1R: 5’-GGGAGTTGTAGTTGTACTCCAGCTT-3’. Transgenic tomato plants were self-pollinated and T_1_ seeds extracted from fruits harvested from each individual line. The T_1_ seeds were germinated on MS basal medium containing 1 mg/L Phosphinotricin, and seedlings that survived under the herbicide selection were transferred to pots containing sterile soil and maintained in a glasshouse at 28-29°C and 80-90% relative humidity. Transgene insertion was confirmed by gene-specific PCR and gene expression confirmed by RT-PCR using the TYLCV *C4*- or *Gfp*-specific primers as described above. For the internal control, a pair of primers for the actin gene (forward primer KL17-071 03g078400F: 5’-TTGCTGGTCGTGACCTTACT-3’ and reverse primer KL17-072 03g078400R: 5’-TGCTCCTAGCGGTTTCAAGT-3’) was used.

### 5.3. Plant RNA Extraction

Total RNA was extracted using 500 mg freshly collected leaf tissue from top third developed leaves of the TYLCV *C4*-transgenic tomato plants (line ‘C4-C1’) in the T_1_ generation as well as from those *Gfp*-transgenic tomato plants as a control, which were in the same developmental stage and growing under the same greenhouse conditions. Each individual leaf tissue sample was processed in a plastic extraction bag using a HOMEX 6 homogenizer (BioReba, Swizerland) with 2.25 ml of TRIzol reagent following the manufacturer’s protocol (Thermo Fisher Scientific, USA). Concentration of the resulting RNA preparation was measured with a NanoDrop micro-volume spectrophotometer (Thermo Fisher Scientific, USA). The quality of cleaned DNA-free RNA preparations was checked in a 1X bleach gel [55].

### 5.4. RNA-Seq library Preparation, Sequencing and Data Analysis

RNA-Seq libraries were constructed as previously described [56]. Six separate RNA-seq libraries were prepared using total RNA preparations extracted from three individual transgenic *C4* plants (T_1_ generation) and three transgenic *Gfp* plants (T_1_ generation). These T_1_ seedlings were 21 days post germination and grown in the same greenhouse with the same environmental conditions of 28-29 °C, 80-90% relative humidity, and 14 h natural sunlight. RNA-Seq libraries were sequenced on an Illumina HiSeq 2500 system to generate 100-bp single-end reads. Adapter trimming and removal of low-quality reads were performed using Trimmomatic [57]. RNA-Seq reads were filtered to remove reads aligned to the ribosomal RNA database [58] using Bowtie [59]. The resulting high-quality cleaned reads were aligned to the tomato reference genome (version SL3.0, The Tomato Genome Consortium, 2012 [60]) using HISAT [61]. Reads were counted for each tomato gene model and normalized to reads per kilobase of exon model per million mapped reads (RPKM). Raw read counts were used as input to the DESeq package [62] to identify differentially expressed genes between the *C4*-transgenic and the control *Gfp*-transgenic plants. Genes with adjusted p-values less than 0.05 and log2fold changes greater than or equal to 1.5 were considered to be differentially expressed.

The Gene Ontology (GO) enrichment analysis of differentially expressed genes was performed using the agriGO program [63]. The Tomato Functional Genomics Database [64] and the iTAK database [65] were used for identification of tomato transcription factors, receptor-like kinases, and microRNA targets. Standalone BLAST [66] was used to identify other genes of interest by comparing them with *Arabidopsis* homologs in conjunction with utilizing annotated GO terms of tomato genes [67].

### 5.5. Validation of differentially expressed genes by qRT-PCR

To validate the differential gene expression as observed in the RNA-seq libraries, 14 DEGs were randomly selected for testing using qRT-PCR. Primers were designed (Supplementary Dataset S5) and their specificity confirmed by aligning the primer sequences to the tomato genome. cDNA was generated from 2 μg of the same tomato RNA preparations as those used for RNA-seq using the SuperScript III cDNA Synthesis System (ThermoFisher Scientific, USA). Twenty-five microliter PCR reactions consisted of 2 μL of diluted cDNA, 0.75 μL of each primer (10 μM), 12.5 μL of 2x Brilliant II SYBR Green Master Mix with low ROX (Agilent), and 9.3 μL of nuclease-free water. PCR amplifications were performed in an Mx3005P Real-Time PCR System (Agilent, USA) using the following cycling conditions: 95°C for 10 minutes, followed by 40 cycles of 95°C for 30 seconds and 60°C for 1 minute with SYBR Green detection during the 60°C step. The presence of a single amplicon in PCR reactions was confirmed by the presence of a single, uniform peak on dissociation curves conducted after amplification. Each of the selected genes was amplified from 3 biological replicates per treatment, with 3-4 technical replicates per biological replicate. Expression levels were normalized to the tomato actin gene (Solyc04g011500) using the ΔΔCt method and expressed in terms of log_2_(fold change) for comparison with the RNA-seq data. Significant differences in gene expression via qRT-PCR was determined using a one-tailed unpaired Student’s *t*-test (if data are normal and homoscedastic), Welch’s t-test (if heteroscedastic) or the Mann-Whitney Wilcox test (if not normally distributed). Statistical analysis was conducted in R (R Core Team 2018 [68]).

## Data Availability

Raw RNA-Seq reads have been deposited in the NCBI SRA database under the accession No. SRP266228.

## Author Contributions

Conceptualization, K.S.L.; methodology, C.P., Y.Z., M.S., J.R.W., Z.F., K.S.L.; validation, J.R.W.; formal analysis, C.P., Y.Z., M.S., J.R.W.; investigation, C.P., Y.Z., M.S., J.R.W.; Resources, K.S.L., Z.F.; data curation: C.P., Z.F., M.S. Writing-original draft preparation, C.P., K.S.L. Writing-review and editing, M.S., J.R.W., Z.F.; Visualization, C.P., K.S.L.; Supervision: K.S.L.; All authors have read and agreed to the published version of the manuscript.

## Acknowledgements

We thank Andrea Gilliard, Deanna Dong, Louis William and Tyler Devaney for their excellent technical assistance, and Bidisha Chanda for reviewing the manuscript.

## Supplementary Materials

**Supplementary Table S1.** Reads summary for the RNA-Seq libraries.

**Supplementary Table S2.** Pearson correlation coefficient among replicate libraries indicate the reproducibility of RNA-seq libraries.

**Supplementary Figure S1.** Original gel pictures used for Figure 2.

**Supplementary dataset S1.** Differentially expressed genes (DEGs) between the TYLCV-*C4* transgenic line (C4-C1) and the control transgenic *Gfp* line (GFP1).

**Supplementary dataset S2.** Up-regulated differentially expressed genes (DEGs) between the TYLCV-*C4* transgenic line (C4-C1) and the control transgenic *Gfp* line (GFP1).

**Supplementary dataset S3.** Down-regulated differentially expressed genes (DEGs) between the TYLCV-*C4* transgenic line (C4-C1) and the control transgenic *Gfp* line (GFP1).

**Supplementary dataset S4.** Pathway analysis of differentially expressed genes (DEGs) between the TYLCV-*C4* transgenic line (C4-C1) and the control transgenic *Gfp* line (GFP1).

**Supplementary Dataset S5.** Quantitative RT-PCR validation of select differentially expressed genes and associated primers.

